# Rational evolution of a recombinant DNA polymerase for efficient incorporation of unnatural nucleotides by dual-site boosting

**DOI:** 10.1101/2022.02.27.482192

**Authors:** Ruyin Cao, Lili Zhai, Qingqing Xie, Zi Wang, Yue Zheng, Wenwei Zhang, Alexander Kai Bull, Xun Xu, Yuliang Dong, Chongjun Xu, Wenping Lyu

**Author notes:** Correspondence: *R.C., X.X., Y.D., C.X., W.L. Warshel Institute for Computational Biology, School of Life and Health Sciences, The Chinese University of Hong Kong (Shenzhen), Shenzhen, Guangdong 518172, P. R. China & School of Chemistry and Materials Science, University of Science and Technology of China, Hefei, Anhui 230026, P. R. China. These authors contributed equally to this work (Ruyin Cao, Lili Zhai).

## Abstract

Machine learning modelling assisting function-oriented enzyme engineering is normally built on predefined protein sequence space. However, efficient defining the determinant amino acid positions upon which the combinatorial mutation library is constructed is still a challenge in protein science. Herein, we present a comprehensive investigation of modifying a recombinant DNA polymerase for efficient incorporating one unnatural nucleotide, including the identification of key sites/regions, machine learning-assisted mutants screening, and the underlying mechanism of kinetics boosting. By using hundreds of training points and only dozens of testing samples, we found that one highly engineered enzyme’s catalytic efficiency can be further improved by one order of magnitude by specific mutation on two sites, 485I and 451L. Compared to the position 485 which is known to dominate local conformation of B-family DNA polymerases, 451 is a split-new active site discovered by our approach. A novel allosteric regulation mechanism is underlying the apparent synergy of 485I and 451L on the kinetics boosting. As a result, a “half-closed” conformation of the binding pocket and a cooperative binding of both primer and template DNA strands on the protein accelerated the processes of substrate’s incorporation, molecular recognition, and releasing of incorrect nucleotides. These findings have implications in guiding the function-tuning of DNA polymerases for a broad range of biotechnological applications.

---

Machine learning (ML) models have been proposed to assist traditional protein engineering protocols in efficiently identifying improved protein variants by reducing the experimental burden of testing many variants [1]. However, ML approaches alone do not offer any mechanistic insights as to how amino acid (a.a.) sequence positions of an enzyme individually contribute to its comprehensive function [2–4]. Defining the determinant a.a. positions upon which the combinatorial space for a specific functional goal is constructed is still a challenge in protein science [5–7]. Practically this aspect is also the key to minimize economical cost in training data collection [1]. A common strategy is to consider potential active sites, such as conserved a.a. positions or those involved in the binding pocket/interface. These approaches might be beneficial in the case of proteins targeting small ligands and those with evolutionarily conserved functional residues [8]. For protein machineries whose active-site residues are highly mutable [9] and function-controlling positions are distant from the catalytic center [10, 11], efficiently identifying key positions still remains challenging [12, 13].

An important example of this challenging category is B-family DNA polymerases [14], the most widespread polymerases in all domains of life exhibiting multiple functional states during nucleotide ligation [15–18]. In modern biotech and biomedicine industries, such as DNA sequencing [19] and next-generation therapeutics [20], the function-oriented modifications of B-family DNA polymerases are highly demanded. In the next generation sequencing (NGS) platforms DNBSEQ^TM^ [21] and COOLMPS [22], Thermococcus kodakaraenis (KOD) DNA polymerase [15], a member of B-family DNA polymerases, has been highly engineered to incorporate synthetic nucleotides modified with cleavable terminator and fluorescent dyes [21, 22]. Notably, the sequencing performance of NGS sequencers is fundamentally driven by the modified nucleotides and paired DNA polymerases. A KOD enzyme with improved one-cycle incorporation efficiency for such nucleotides would greatly advance accurate sequencing by synthesis chemistry-based NGS.

Aimed at enhancing the catalytic performance of a highly engineered parent KOD incorporating a specifically-modified nucleotide, we present here a comprehensive investigation from the identification of key sites/regions, to ML-assisted mutant screening, to the underlying mechanism of boosted kinetics. After *in silico* functional dynamics analyses of the KOD at atomistic level, a medium-sized dataset (349 samples covering 70 sites) was collected to construct our ML models. The catalytic performance of the predicted KOD mutants (33 samples) achieved as high as 31 folds relative to the parent KOD with only 10% experimental costs of the training data. Importantly, the best-performed KOD mutant bears double mutations outside the polymerase’s active center, which are unreachable by general-purpose active-site prediction strategies. By diving into the synergetic effect of the two mutation sites on the boosted kinetics, we discovered that stabilizing the DNA duplex structure *via* specific combination of far-end a.a. 485 and 451 could accelerate the incorporation efficiency of substrate with a “half-closed” binding pocket, implying an unexplored regulation mechanism of the polymerase functionality.

## Structural insights into the functional dynamics of KOD

KOD adopts a disk-shaped structure comprising five subdomains [17], i.e. the N-terminal (N-term.), exonuclease (Exo.), palm, finger, and thumb (Figure 1A). The DNA duplex (PT) composed of one primer and one template is bound within a groove jointly formed by the palm and thumb, while incoming nucleotide is added to the 3’-OH end of the primer at the catalytic cavity jointly formed by finger, palm and thumb. The incorporation process of natural nucleotide involves significant polymerase conformational changes in five steps (Figure 1A) [10]. Firstly, binding with PT induces the thumb subdomain to transition from a partially disordered apo state to a well-ordered binary state (state I) [17]. As nucleotide is loosely accommodated into the active pocket (state II), the finger subdomain undergoes an inward tilt toward the palm domain, leaving the nucleotide tightly trapped in a precatalytic state (state III) [17]. Next, the ternary complex experiences nucleotidyl-transfer [23], pyrophosphate (PPi) dissociation (state IV) and finger reopening (state V) [10]. In NGS application, polymerase translocation along PT (state V back to state I, Figure 1A) is blocked by the incorporated modified nucleotide as it carries a 3’-OH terminator [21, 22]. Therefore, conformational changes of the polymerase are the rate-limiting steps in one catalytic cycle of single-nucleotide incorporation [10].

**Fig. 1.**
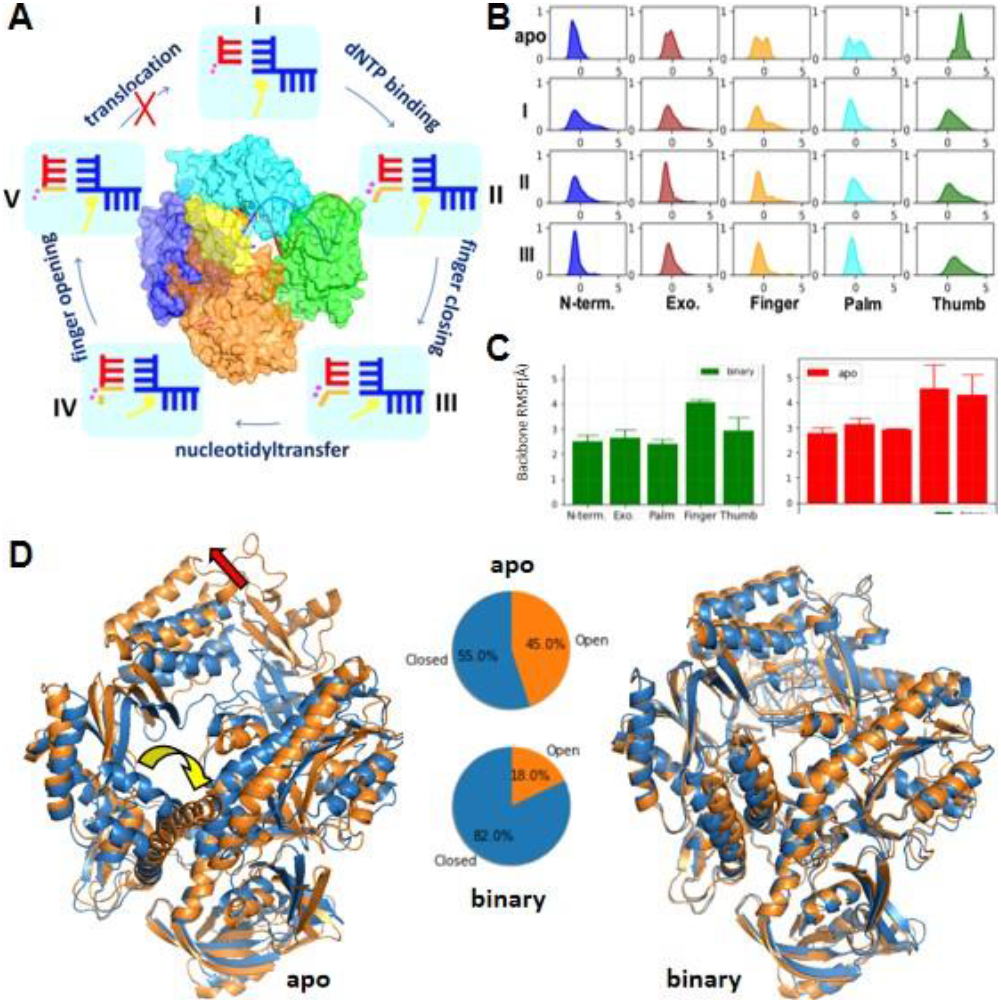
Conformational dynamics of KOD enzymes. **A)** Functional subdomains of KOD and kinetic scheme of its incorporation cycle. The subdomains of KOD are rendered by different color codes: N-terminal (blue), exonuclease active site (sand), finger (yellow), palm (cyan) and thumb (green). Starting from a DNA-bound binary state (I) with open finger (yellow) and unoccupied active site, one nucleotide enters the active site (II) and the finger domain closes to establish tight binding with incoming nucleotide; the nucleotide is then added to the 3’-end of primer strand and pyrophosphate is released from the active site (IV); note that the translocation of DNA double strands can be hampered by a 3’-terminated nucleotide. **B)** Change of KOD backbone fluctuations across apo state (1WNS), binary state (I, 4K8Z), finger-open ternary state (II, 5VU7), and finger-close ternary state (III, 5OMF), characterized by the density distribution of Ca atoms’ normalized B-factor. **C)** The statistics of RMSF of each functional subdomain of the parent KOD (RF) in apo and binary states. **D)** The most weighted (blue) and the second weighted (orange) representative conformations of the RF sampled in apo and binary states. The yellow and red arrows highlighted the local conformational transforms between the two representatives in apo. The probability of ‘open’ and ‘closed’ states were estimated by the weightings of the representatives in clustering analyses

In addition to the apo state [15], KOD complex states I (binary) [16], II (ternary with open finger [17]), III (ternary with closed finger [18]) have been captured by X-ray crystallography. Via analyzing B’-factor distribution [24] of *C_α_* atoms deposited in the X-ray data, we noticed that the atomic fluctuations of the exonuclease, finger, palm, and particularly the thumb subdomains are all notably suppressed in these complex states (I-III, Figure 1B). We then performed μs-scale all-atom molecular dynamics (MD) simulations to in detail assess structural dynamics of one BGI-patented recombinant KOD (WO2018CN103764, named as RF). Here, we focused on modelling RF in apo state and state I. On one hand, the finger and thumb in apo RF simulation show the greatest flexibility, in line with static structure-based insights (Figure 1C). Particularly, tip of the finger subdomain (fingertip), responsible for recognizing the incoming nucleoside [10, 25], is observed to swing between the palm and exonuclease subdomains (Figure 1D), thus inter-inducing the “closed” and “open” conformations of the enzyme. More notably, the inward movement of the fingertip from palm to exonuclease subdomains (yellow arrow, Figure 1D) coincides the outward movement of the thumb region (red arrow, Figure 1D), resulting in a looser binding pocket [26] of RF for PT. On the other hand, backbone flexibility of this polymerase is greatly suppressed in the binary complex state (Figure 1C), with thumb being the most suppressed. Despite still being the most flexible, the finger subdomain exhibited a higher probability (55% in apo state vs. 82% in binary complex, Figure 1D) of adopting the “closed” conformation (the fingertip swinging toward palm region) in binary complex. The “close” finger is believed to be necessary to improve the catalytic efficiency [27, 28]. Yet, the closed-biased conformational distribution might also limit the kinetics of state transitions of I→II and IV→V in the catalytic cycle (Figure 1A).

## Machine learning-assisted mutant screening towards boosted kinetics

The overall kinetic performance of one KOD mutant can be described by the Michaelis-Menten kinetic parameter *V_max_/K_m_* [29, 30]. Then, the catalytic efficiency of KOD mutants is characterized by the ratio of *V_max_/K_m_*\*mut* relative to that of the RF:

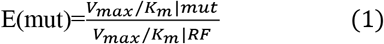

Based on the structural insights of crystallized KODs and the RF (Figure 1), we built a ML-assisted mutant screening protocol in 3 steps (Figure 2A): 1) preliminary screening of mutations on all the responsive regions upon the binding of PT (namely the finger, thumb, palm, and exonuclease subdomains) and mutation effectiveness-to-mutation region analysis; 2) ML models design and training; 3) unexplored mutants’ prediction and experimental validation.

**Fig. 2.**
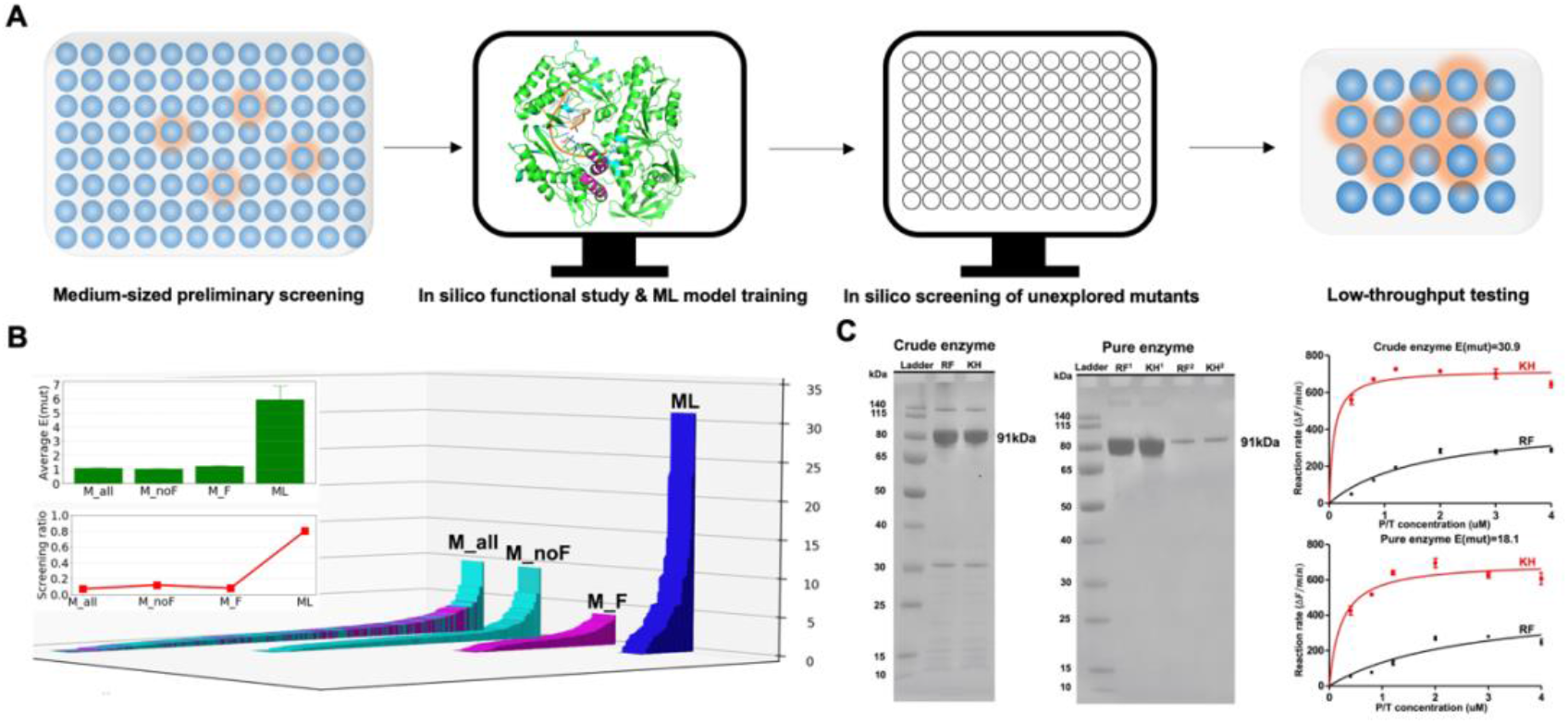
Machine learning-aided mutant screening of KOD. A) Scheme of Knowledge-based ML screening workflow of KOD. Note residues involved in preliminary screening (colored in cyan) and virtual library design (colored in purple) are highlighted in the 3-D structure of the parent enzyme RF. B) Statistical distributions of tested E(mut) values of KOD mutants in preliminary screening step (M_all: all tested mutants, M_noF: mutations on non-finger regions; M_F: mutations on finger region) and machine learning-accelerated step (ML). The illustrations are the average M(mut) and screening ratio (coverage of the tested mutants to the full combinatory space of sequences) of the M_all, M_noF, M_F, and ML subsets, respectively. C) SDS-PAGE analysis of crude enzymes (left panel) and pure enzymes (right panel) of RH and KH and Kinetic efficiency comparison between RF and KH for single-step incorporation of one dATP analogue. Note in the pure enzyme, RF^1^ and KH^1^ denote 5mg enzymes while RF^2^ and KH^2^ represent 0.25 mg enzymes.

Traditional methods including random mutagenesis [31], site-directed mutagenesis [32], and semi-rational design [33] were utilized in preliminary screening of mutations on finger, thumb, palm, and exonuclease subdomains. Here, positions in the active site were fixed, as the RF has been highly engineered towards incorporating BGI tailor-made nucleotides [21, 22] and the chemistry step of nucleotide insertion itself is not the rate-limiting step in our application [10, 21, 22]. At the expense of quantifying single-nucleotide incorporation kinetics for 349 crude KOD mutant enzymes covering 70 positions (noted as M_all), one optimal mutant exhibiting remarkable kinetic enhancement (E(mut) ~9) and a few less optimal mutants (E(mut) > 4) (Figure 2C) were discovered.

However, the average E(mut) of all these mutants is around 1 (upper inset, Figure 2C), implying that the preliminary engineering resembles a random walk in the fitness landscape of E(mut). One possible reason for difficult lifting of E(mut) is that the 349 tested mutants are just a very small subset (<2%, upper bound, Figure 2C) of the combinatorial space covering hundreds of a.a. sites. Also, it could be that the E(mut) landscape of KOD is essentially epistatic enriched with low-fitness “functional holes” [34].

Interestingly, KOD variants involving mutations of the finger subdomain (121 mutants covering 39 positions, noted as M_F, upper inset of Figure C) present a discernible improvement of E(mut) (averaged at 1.22±0.07) compared with the other finger-exclusive mutants (228 mutants covering 44 positions, noted as M_noF) (averaged at 1.03±0.07), albeit the screening ratio (upper bound) of M_F (<3%) is only 1/5 of that of the M_noF (14%, upper bound). These experimental results indicate that the mutations on the finger subdomain are effective ones toward increasing E(mut), consistent with our insights obtained from MD simulations. Nevertheless, the combinatorial complexity of mutations on the whole finger subdomain (covering 52 a.a. sites) is still a great challenge for experimental kinetic evaluation.

ML models were then employed to rationally identify undiscovered mutations on the finger subdomain towards lifting E(mut) at low experimental throughputs. Firstly, hidden patterns between mutations and kinetics within the tested pool of KOD mutants were learned by 3 extra tree regressor models [35] on 24 sequence-based and structure-based features. Then, based on structural insights into the functional dynamics of RF and the existing pool of KOD mutants’, kinetics data, we designed a virtual variant library covering six finger-located positions: a.a. 451, 453, 480, 484, 485, 486 (purple color representation in Figure 2B). In the preliminary screening, single mutation on a.a. 451, 453, 480, 485 exhibited moderate boosting effects on enzyme kinetics (1.85-3.90 folds). The neighbors of a.a. 485, a.a. 484 and 486 were also considered. Note, all the untested variants included in this virtual library contains one mutation on a.a. 485 with respect to RF. This site has also been engineered in another unnatural KOD targeting incorporation of xeno-nucleic acid (XNA) [17]. In total 159 virtual variants were assessed by the 3 regressors, and the 33 top ranking mutants (scored by the mean value of the predicted E(mut) from 3 ML models) were experimentally validated.

For each of the 33 ML-predicted mutants, we measured the E(mut) (labeled as ML in Figure 2C), using the same experimental protocol as in the preliminary screening step. Out of this small testing pool, 31 variants showed enhanced kinetics in the experiment (E(mut) >1, true positive rate 94%). Particularly, 6 variants with double mutations on a.a. 451, 484 and 485 showed E >10, among which one variant KH reached the highest E=31.5 (Figure 2C). Compared to the best performed dual-site mutant on the finger region without the assistance of ML (E(mut)=4), KH saw improvement in E(mut) by around 7 folds with around 1/3 of the former’s experimental costs, indicating this ML-assisted strategy exerted on the finger subdomain greatly improves the screening efficiency.

Note that the KH only differs in the two single point mutations S451L and E485I from its parent RF. We further reevaluated the kinetic performances of both the KH and RF at higher degree of enzyme purity (right panel of Figure 2D, 90% purity). Although the activity of enzymes might be harmed in the purification process, the KH still exhibited 18 folds rate acceleration (Figure 2E). We conclude that dual-site mutations on the finger subdomain can enhance the polymerase’s kinetical performance by at least one order of magnitude.

## Mechanism underlying the dual-site boosting

It is worth noting that single mutations on a.a. 485 or 451 of the RF achieved only 2 folds improvement of the catalytic efficiency, impressively underperforming their combined mutations introduced to the KH (18 folds). In other words, it is the synergy of S451L and E485I that contributes to KOD’s comprehensive kinetic improvement. However, the two a.a. sites are evolutionarily uncorrelated (Figure 3A) and their respective degrees of conservation [36] are distinctly different (a.a. 485 is highly conserved to ala-nine whereas a.a. 451 is non-conserved, Figure 3B). Consistent with their weak evolutionary correlation, the two sites are not in spatial proximity (*C_β_-C_β_* distance in crystal: 21.6 Å, PDBID:5omf).

**Fig. 3.**
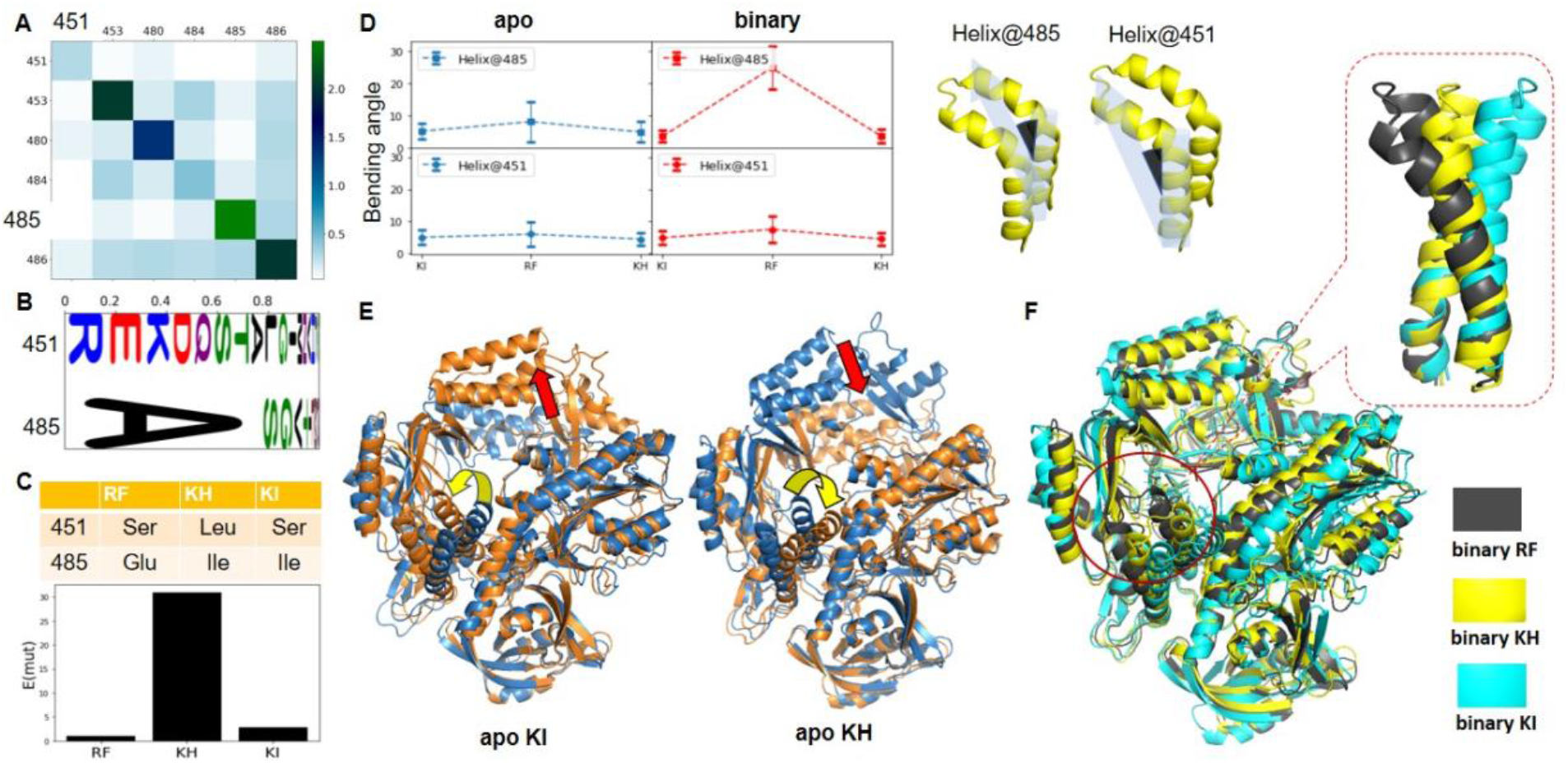
Sequential and structural comparison of the parent KOD RF and ML predicted KOD mutants. A) SCA-based evolutionary covariance score matrix of KOD on a.a. 451, 453, 480, 481, 485, 486. Higher score means stronger evolutionary correlation between pairwise positions. The correlation between a.a. 451 and a.a. 485 is around 0.04. B) Probability-based sequence logo of KOD in a.a. 451 and 485. C) The type of mutations on the a.a. 451 and 485 sites of the RF, KH, and KI (upper panel); the measured E of the three mutants (lower panel). D) The bending of the finger subdomain of the RF, KI, and KH in MD simulations are characterized by the bending angle of the two finger helices (Helix@485 and Helix@451). Right panel represents the definition of the two angles. E) the most weighted (blue) and the second weighted (orange) representatives of the KI and KH sampled in apo state, respectively. The yellow and red arrows highlighted the local conformational transforms between the two representatives. F) Superposition of the most weighted representative of the RF, KH, and KI in binary state. The enlarged plot highlights the conformational difference on their finger subdomains.

We then explored the synergy of the two sites on the structural dynamics of KH via MD simulations. Similar to the case of RF (Figure 1), we performed three independent all-atom MD simulations for the KH in both the apo state and the binary complex state (6 μs in total). For the purpose of comparison, MD simulations of apo and binary states at the same length scale were also carried out for a single-site mutant bearing only 485I (noted as KI, E(mut)=2.6, Figure 3C). Firstly, we analyzed the conformational change of the two helices of the finger subdomain where the two mutations are located (Helix@451 and Helix@485, Figure 3D). Results show that the average conformations of the finger subdomain are indistinguishable among the KH, RF and KI in apo state (Figure 3D). In fact, the fingertips of the three systems are “free” to adopt the open and/or closed conformations regardless of the two sites’ a.a. compositions (Figure 3E and Figure 1E). After the binding of PT, the mutation E485I leads to conformational differences of the finger subdomain significantly (Figure 3D): the helix@485 in both the KH and KI are less bent towards the palm region than in their RF counterpart (reduced from 20° to 5° on average, Figure 3D). The bending angles of helix@451 of the KH and KI are also slightly decreased. Overall, the finger subdomain of the two mutants is more likely to avoid the closed conformation than that of the parent RF in the binary complex state (Figure 3F), resulting in a larger space for the binding of single-nucleotide in the functional states I-III (Figure 1B). This increase in available space might be beneficial for the incorporation of a bigger nucleotide attached with the reversible 3’-O blocking group and a fluorescent dye, thus improving the kinetical performance. However, the local conformational shifting of the finger domain is indistinguishable between the KH bearing the double mutations and the KI carrying only single mutation on a.a. 485 (Figure 3D). We thus conclude that the local conformational shifting of the finger relies mainly on the mutation on a.a. 485 and it becomes significant after the binding of PT.

There is no direct interaction between sites 451, 485 and the nucleotides of PT duplex, as the two residues are located at the back of the PT binding groove (Figure 4A). The communication (if any) between the two sites and PT must be mediated by other subdomains. Several residues in close proximity to a.a. 451 and 485 (closest heavy atom distance less than 0.5 nm on average) are detected in the N-term. (a.a. 364), palm (a.a. 446), and Exo. (a.a. 329, 332, 333, and 336) subdomains (Figure 4B-C). Interestingly, the KH and KI share the same mutation on a.a. 485, while two close contacts (485—329 and 485—332) are missing on the KI (Figure 4B). On the other hand, the two close contacts of a.a. 451 (451—364 and 451—446) are both indistinguishable with respect to their average distances for all the three KODs (Figure 4B), although KH is mutated from KI and RF at a.a. 451.

**Fig. 4.**
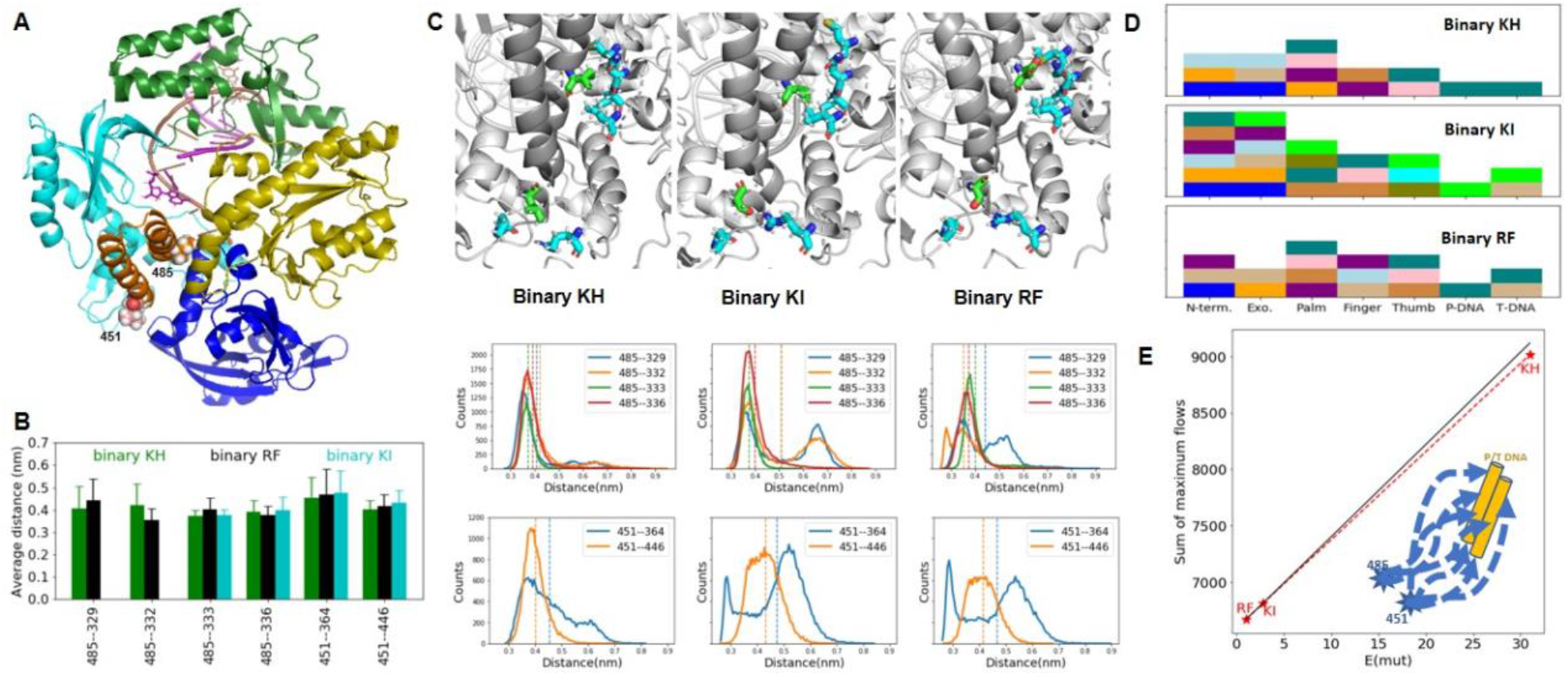
Conformational dynamics analysis of the KH, KI, and RF in binary state. A) the relative positions of the two mutation sites and the P/T DNA in 3-D structure; B-C) inter-subdomain close contacts of a.a. 451 and 485 of the three systems. The average distances and their distributions are reported in B) and lower panel of C), respectively. The representative conformation of the close contacted residues is showed as illustration for each system. The vertical dashed lines marked the average distances, respectively. D) a clustering characterization of the dynamical correlation of residues/nucleotides of each subdomain and P/T DNA of the KH, KI, and RF, respectively. Here, different color code represents different cluster ID of each system, while two subdomains sharing a same color means their conformational dynamics are correlated. E) the correlation between the measured E and the sum of maxima flows of correlated movements from the two mutation sites (a.a. 451 and 485) to the P/T DNA. The black solid line represents an ideal correlation estimated from RF and KI only. A schematic diagram of the maxima flows from the two sites to the P/T DNA is presented as illustration in the lower side.

To assess detailed differences in protein dynamics resulting from distributions. For the single-site mutated KI and the parent RF, dual-peak distributions are observed for one (451—364) and three (451—364, 485—332 and 485-329) inter-subdomain contacts, respectively. These results indicate that the mutations are more consistent with a change of the dynamics of the residue-to contacts, rather than with changes of the average structures.

We further characterized the pairwise dynamical correlation of all residues and/or nucleotides by the normalized covariance of their center-of-mass, and then learned the dynamical cooperative residues and/or nucleotides through a parameterless unsupervised classification method [37, 38]. Results show that the dynamics of DNA duplex is not directly correlated to that of the finger subdomain in all the three KODs (they are assigned to different groups, Figure 4D), but partially correlated to the Thumb, Palm, and Exo. subdomains. Interestingly, the entire PT-DNA duplex is clustered in a single cooperative group only in the binary KH (Figure 4D). For the RF and KI, the T-DNA strand is split into two correlation groups around the 5’-terminal where only a correct incoming nucleotide should be base-paired. As an extended correlated dynamical network around reaction center could enhance catalysis of an enzyme [39], the core of a binding pocket (the 5’-terminal nucleotide of T-DNA) linked to an internally-cooperative duplex DNA structure might also benefit base-pairing, implying another possible mechanism of the improved E of KH.

Finally, we quantified the correlation between different mutations and the structural dynamics of PT duplex, by estimating the sum of maximum flow [40] of correlated movements (∑*MaxF*) stemming from a.a. 451 and 485 to each nucleotide of PT mediated by all possible residues/nucleotides (illustrated as dot line arrows in the lower side of Figure 4E). Higher the ∑*MaxF*, greater the capacity of information (in the form of cooperative conformational dynamics) spread from the two sites to the DNA duplex (and vice versa). Results show that the ∑*MaxF* is surprisingly well correlated to the E(mut) values for the RF, KH, and KI (red dashed line, Figure 4E). The best performed KH has the highest communication capacity of conformational dynamics between the dual-site mutations and PT.

Taken together, we conclude that the observed enhancement of incorporation kinetics presented by KH is a combined result of changes in conformational dynamics within local enzyme subdomains and the concerted dynamical motion of the bound DNA duplex linked to the core of binding pocket.

## Conclusions

We successfully enhanced the catalytic efficiency (more than one order of magnitude) of a highly engineered KOD enzyme towards non-natural nucleotides for better performance in sequencing applications. The dual-site mutations (451L and 485I) of the best performed KH show a significant cooperative effect on the enhancement of kinetics. Namely, 485I regulates the local conformation of the finger subdomain, which dominates the processes of “capture” and “release” of single nucleotide; while 451L modulates the internal dynamical cooperation of the PT duplex, which extends the correlation network around the core of the binding pocket. It is worth to remind that no co-evolution in sequence between the newly discovered a.a. 451 and the active site 485 reported elsewhere. In particular, the a.a. 451 itself is a non-conserved site and the effect of mutations on the average conformation of the finger subdomain is insignificant. These properties imply that the site 451 could be easily ignored by traditional active-site prediction protocols. Therefore, we propose that rational integration of detailed structural insights and knowledge-based machine learning models, is a powerful strategy to efficiently accelerate the engineering of a complex enzyme towards a targeted biological function. We believe that both the strategy and the findings presented in this work will enable advanced protein engineering of other polymerases or other nucleotide-involving protein machineries in the future.

## Materials and Methods

### A. Reagents and instruments

DNA templates and primers labeled at the 5’-terminus by CY5 dye were synthesized from Invitrogen. 200μL of Labeled primer(100μM) was annealed to 200μL of the DNA template (100μM) by heating to 80°C for 10 minutes in thermomixer and then slowly cooling down to room temperature by turning off the power. Mutant primers were purchased from GenScript, and dNTPs, Pfu DNA polymerase and Dpn1 were bought from NEB. Modified 3’-blocked dATP (dATP-CY3) by Cy3 dye was supplied by BGI synthetic chemistry group. Sequences of parent enzyme RF and all mutants were integrated in a commercial vector pD441-pelB from DNA2.0. LB broth, kanamycin and IPTG were bought from Thermo Fisher. Affinity chromatography column (His Trap HP 5ml, GE), ion exchange column (HiPrep Q FF, 5mL and HiTrap SP HP, GE) and AKTA were purchased from GE. Protein concentration was measured by using Bradford method with a microplate reader from TECAN. PCR reactions performed in a Bio-Rad LifeECO™ Gradient Thermal Cycler were carried out in a Cycler 96 system with visualization of DNA for motion via SYBR safe (Invitrogen).

### B. Library construction and protein expression

Site-mutagenesis strategy was performed for library construction. The PCR reaction components consisted of 10xPfu Buffer with MgSO4 (2.5μL), 10mM dNTPs mix (0.5μL), DNA template (75ng), Pfu DNA polymerase (1U), nuclease free water to 25uL. Thermal cycling conditions were 95°C for 3 minutes denaturation step, followed by 20 cycles at 95°C for 30 seconds, 53°C for 30 seconds, 72°C for 7 minutes and final extension at 72°C for 10 minutes. Then, 1μL of DpnI was added to each PCR tube to digest template DNA at 37°C for 1 hour. 5μL of PCR product of each mutant was transformed into the E. coli DH5α competent cell. Preparations of recombinant plasmids were conducted using the QIAprep Spin Miniprep Kit from QIAGEN, and then these plasmids were sequenced by BGI Group (Genome Sequencing Company). Next, those recombinant plasmids with the expected sequencing results were transformed in E. coli BL21 (DE3), and then one colony was selected and cultivated in 500μL LB medium containing 50ug/ml kanamycin using 96-deep well plates as the first-class seed. Second generation was cultivated in 50mL of LB medium in 250mL shaking flask and introduced by IPTG with final concentration of 0.5mM to express overnight at 25°C in plate shaking incubator.

### C. Crude Enzyme screening

Polymerase reaction started by adding DNA polymerase to a premixed solution containing DNA primer-template complex attached with CY5 (PT, Scheme S1), modified dATP attached with 3’-blocked terminator and CY3 (dA’, Scheme S1), BSA and screening buffer (20mM Tris-HCl, 10mM KCl, 10mM (NH4)2SO4, 0.1% Triton, 4mM MgSO4, pH 8.8) at 40 °C. Reaction progress was monitored via fluorescence resonance energy transfer (FRET) signals from Cy3 (excitation at 530nm) to Cy5 (emission at 676nm)) over time using the Spark Control plate reader (TECAN) in a corning 384 black plate (clear bottom) [41]. Polymerase reactions contained 0.1μM PT, 0.25μM dA’, 2μg BSA, 1μg of DNA polymerase (crude enzyme) and nuclease free water to final volume of 50μL. RF was used as the positive control, and enzyme replaced by screening buffer was used as negative control. Crude enzyme PMSF and 1mg/ml lysozyme, pH 7.6) with a ratio of 0.04g pallet cell per mL buffer. The resuspended cells were incubated at 37°C for 10 minutes, followed by thermal denaturation at 80 °C for 30 minutes. After centrifugation at 12,000xg for 20 minutes, the supernatant was collected, and the crude enzyme concentration was determined. The purity of these crude enzymes was checked via SDS-PAGE, and then directly used for the enzyme activity screening.

### D. Purified enzyme preparation

Purified enzymes of RF and KH were obtained by the following methods. Pellet precipitation was suspended in lysis buffer 2 (500mM NaCl, 5% glycerol, 20mM imidazole, 1.25mM PMSF, 50mM potassium phosphate, pH7.4) and then crushed with a homogenizer. After centrifugation, the supernatant was filtered with 0.22μm pore size membrane. Next, 10mL of the sample was loaded to a HisTrap FF column (5ml) pre-equilibrated with 25mL buffer 2, and then washed with 50mL buffer2, followed by 50mL elution buffer (500mM NaCl, 5% glycerol, 500mM imidazole, 50 mM potassium phosphate, pH 7.4) at speed of 3mL/min. The eluate was loaded to a HisTrap Q HP column (5ml), and then eluted with a buffer containing 5% glycerol, 25mM potassium phosphate, pH6.6, and the flow through was collected. Furthermore, the collection liquid was upload in a HisTrap SP HP (5ml), and then linear eluted by 0%→60% buffer 3 (1M NaCl, 5% Glycerol, 50mM Potassium Phosphate, pH 7.4) and the first peak was collected. The eluate was dialyzed with buffer 4 (20mM Tris 200mM KCl, 0.2mM EDTA, 5% glycerol, pH 7.4) for 18 hours, and then stored in 50% glycerol in final concentration 1mg/mL. Purified enzymes were stored at −80°C until being used. The enzyme purity was at least 90% as determined by SDS-PAGE.

### E. Kinetic measurements of KOD variants

We determined kinetic parameters in terms of Vmax and Km for each enzyme variant using the “Michaelis-Menten equation” aa way of quantifying variants’ incorporation efficiency toward dA, (40). We assume that the Vmax/Km ratio between two variants provide a straightforward appraisal on their comparative catalytic efficiency. In this study, 0.1μM polymerase and 4μM dA’ were used. Reaction rates were tested with varying concentrations of PT (0.4μM, 0.8μM, 1.2μM, 2μM, 3μM, 4μM) and Km and Vmax can be obtained from Michaelis-Menten equation through least-squares fit- ting by GraphPad Prism (30). RF was used as the positive control and its Vmax/Km value was defined as 1, and the E values of all KOD mutants (crude enzymes) were calculated according to eq. 1. Furthermore, using the same kinetic testing methods as for crude enzyme (Reaction rate vs. PT concentration), Vmax and Km of purified RF, KI, KH (90% purity) were also calculated. Besides, Vmax and Km of purified RF, KI, KH were measured with varying dA’ concentration (0.2μM, 0.5μM, 1μM, 2μM, 3μM, 4μM), 0.1μM enzyme and 4μM PT.

### F. Machine learning modelling and extensive MD simulations

The machine learning (ML) models built in this study were aimed at identifying KOD variants with enhanced E values compared to RF. Model building was based on characterized E of a medium-sized pool of KOD variants all seeded from RF. For each kinetically characterized variant, its mutation contents with respect to RF and assay-based E value of crude enzyme was collected for model construction. In one ML model, each variant was represented as a vector encoding sequence-based and structure-based features. The primary sequence of each variant was fed into a ProtFeat python class [42] to generate common protein descriptors, including PI point, flexibility, charge, aliphaticness, total entropy, instability, aromaticity, gravy, mutability scale, volume scale, polarizability scale, ja aascale, asa in tripeptide scale. Apart from including features directly generated from ProtFeat, three features were also calculated to quantify the sequence similarities of whole protein, finger subdomain, thumb subdomain between the variant and RF. Besides, a score derived from PROVEAN [43] was calculated for each variant to quantify the functional effect of site mutations. Structural features of each variant include solvent accessible surface areas (SASAs) of hydrophobic residues located respectively in whole protein, finger and thumb subdomains, backbone B-factor averaged across the whole protein, finger and thumb subdomain as well as the binding free energy be- tween the variant and the DNA double strands. Structural models of apo state and binary complex state for each variant were built with modeler v9.23 [44]. Structural features were calculated based on conformations sampled from MD performed with AMBER18 suite of programs [45]. The machine learning models built here were implemented with the Extremely Randomized Trees algorithm [35] within Scikit-learn [46]. Extremely Randomized tree is one ensemble learning technique which aggregates results from M decision trees to output a mean result for regression problems or a majority vote for classification problems. Each Decision Tree in the Extra Trees Forest is built from the original training input, without random bootstrapping as used by Random Forest. For each decision tree, a random subset of K features is drawn, and selected features are split on random cut points and the best feature chosen for each node is chosen based on the total reduction of some mathematical criteria (typically the Gini Index). The rationale behind the Extra Trees method is that the explicit randomization of the cut point and features combined with ensemble averaging should be able to reduce variance. The dataset used for ML model building has 349 samples in total and was copied into 3 child datasets using Y-stratified train_test_split function (ratio between training samples and testing samples is 7:3) with different random states. For each child dataset, the training subset and testing subset based on Y-stratified splitting were respectively used for regressor building and performance evaluation, and hyperparameter tuning was implemented using 5-fold GridSearchCV. The best model selected from each child dataset was then aggregated to output a mean E value for each variant. One restricted library including 159 variants carrying two mutations on the finger subdomain with respect to RF was constructed based on structural insights into KOD functioning. Each variant was assigned a mean predicted EML score by the three ML models and top 20% variants (33 in total) according to score ranking were then expressed by site-directed mutagenesis and kinetically characterized. For extensive MD simulations with respect to mechanistic study, structural models of RF, KI, KH in apo and binary complex states were constructed with modeler v9.23 [44] based on crystal templates [15, 17]. For each of RF, KI, KH, 3 independent 1-μs MD simulations of both apo state and binary state (18 in total) were performed using AMBER18 suite of programs [45], with the CUDA implementation for GPUs. Protein, DNA du- plex and water atoms were respectively treated with ff14SB, OL15 and TIP3P force field parameters. Each model was solvated in an octahedral water box, with Cl- or Na+ ions added to neutralize the system. Each solvated system was minimized using gradually decreasing restraint on heavy back-bone atoms, heated up to 313K in NVT ensemble and equilibrated for 1 ns in NPT ensemble. Three 1-μs production MD simulations were independently performed for each model.

## Acknowledgements

This work was supported by Shenzhen Engineering Laboratory for Molecular Enzymology (No. DRC-SZ [2018]958). W.L. appreciates National Natural Science Foundation of China Grant No. 21505134.

## Notes

### Competing Interest Statement

The authors have declared no competing interest.

